# Intrarenal Generation of Aldosterone Contributes to Ischemia-Induced Hypertension and Nephropathy in Mice

**DOI:** 10.1101/2025.08.24.671658

**Authors:** Ziwei Fu, Chang-Jiang Zou, Alex Kimball, Tianxin Yang

**Author notes:** Correspondence to: Tianxin Yang, M.D., Ph.D., Division of Nephrology and Hypertension, University of Utah and Veterans Affairs Medical Center, 30N 1900E, RM 4C224, Salt Lake City, UT 84132.

## Abstract

**Background:** The past few years have witnessed a significant advancement in aldosterone (Aldo)-targeted therapies for the management of treatment-resistant hypertension and chronic kidney disease, which often exist in tandem. While Aldo is believed to predominantly originate from the adrenal glands, this study provides evidence to support the involvement of intrarenal Aldo biosynthesis in the pathogenesis of ischemic nephropathy and hypertension in a two-kidney, one-clip (2K1C) model.

**Methods:** We generated inducible renal tubule-specific deletion of C11B2 (RT C11B2 KO) and characterized the phenotype during the 2K1C procedure. We investigated the underlying mechanisms involving the use of mice with collecting duct-specific (pro)renin receptor (CD PRR KO) or renin (CD renin KO).

**Results:** RT C11B2 KO induced a partial blockade of the hypertensive response but a much greater improvement in renal fibrosis and inflammation four weeks post-2K1C. This phenotype was associated with an effective blockade of intrarenal generation of Aldo contrasting to unchanged circulating Aldo concentrations. Furthermore, clipping-induced acute kidney injury was also attenuated 24 hours post-2K1C, at which point blood pressure (BP) did not increase. Similarly, sodium nitroprusside effectively lowered BP in C57BL/6j/2K1C mice but failed to improve renal injury. Additionally, we identified CD) PRR and renin as key upstream regulators of intrarenal Aldo biosynthesis following the 2K1C procedure.

**Conclusions:** Together, these results support the idea that CD PRR/renin-dependent generation of intrarenal Aldo may play a primary role in the pathogenesis of ischemic nephropathy and a secondary role in hypertension development during renovascular constriction. Therefore, targeting intrarenal Aldo biosynthesis may represent a more effective and safer intervention than existing Aldo-targeted therapy to manage ischemic nephropathy as well as hypertension.

## Introduction

Renovascular hypertension, the most common form of secondary hypertension often results from obstruction of the main renal artery due to either atherosclerotic occlusion or fibromuscular dysplasia.^1^ This disease is commonly associated with kidney injury as a consequence of renal ischemia (ischemic nephropathy) or elevation of blood pressure (BP) (hypertensive nephropathy). Activation of renin-angiotensin system (RAS) is well known to play a key role in the pathogenesis of renovascular hypertension, as exemplified by the 2-kidney, 1-clip (2K1C) renovascular hypertension model.^2^ While the increased release of renin from juxtaglomerular (JG) cells and subsequent activation of systemic RAS are well described in the 2K1C model, our recent study reveals an essential role of collecting duct (pro)renin receptor (PRR) and the intrarenal RAS.^3^ Anti-RAS regimen is effective in managing hypertension and cardiorenal disease, but its efficacy is limited particularly when fibrosis is established. ^4^ Furthermore, this therapy sometimes causes an acute decline in renal function due to compromised renal blood flow in the ischemic kidneys.^1^ Several prospective, randomized trials for atherosclerotic disease have failed to demonstrate that renal revascularization is more effective than medical therapy for most patients.^1,5^ Therefore, these studies underscore the need to understand the mechanism of renovascular hypertension and to develop novel therapies for this disease.

Recently, several innovative drugs in clinical trials have focused on aldosterone (Aldo).^6–8^ Double-blind, randomized, placebo-controlled phase 2 clinical trials have consistently shown that the inhibition of Aldo synthase, cytochrome P450 family 11 subfamily B member 2 (C11B2) with Baxdrostat or Lorundrostat leads to a substantial reductions in blood pressure (BP) in patients with treatment-resistant hypertension (TRH)^8^. Both inhibitors have entered Phase III clinical trials. In addition to the BP-lowering effect, Lorundrostat lowered albuminuria by 37–40% in patients with chronic kidney disease^9^. Indeed, excessive Aldo may stimulate renal Na+ reabsorption, expand plasma volume, and elevate BP, it can also promote inflammation and fibrosis in myocardial, vascular, and renal tissues. All of these effects contributes to the progression of TRH and CKD severity.^10,11^ Aldo biosynthesis is believed to occur predominantly in the zona glomerulosa cells of adrenal glands. A number of extra-adrenal tissues such as the brain, blood vessels, the kidneys, and adipose tissues have been reported to produce small quantities of Aldo. However, the functional importance of extra-adrenal Aldo production has not yet been established. In this study, we used the Pax8 system to produce renal tubule-specific deletion of C11B2 (RT C11B2 KO) and examined the phenotype in the 2K1C model. We established the essential role of intrarenal C11B2 in the pathogenesis of clipping-induced nephropathy and hypertension. Furthermore, our findings support intrarenal Aldo biosynthesis as an integrative component of intrarenal renin-angiotensin-aldosterone system (RAAS) under the control of CD PRR.

## Methods

### Animals Care

All animals were cage-housed and maintained in a temperature-controlled room with a 12:12-hour light-dark cycle, with free access to tap water and standard mice chow. All animal studies conducted in the present study were approved by the Animal Care and Use Committee at the University of Utah. All husbandries, handling procedures, animal health monitoring, diet, environmental controls meet the standards described in the Animal Welfare Regulations and the Guide for the Care and Use of Laboratory Animals. The Animal Resources Center at the University of Utah is AAALAC accredited.

### Generation and Genotyping of KO Mice

The C11B2 gene (NCBI Reference Sequence: NM_009991) is located on mouse chromosome 15. It consists of 9 exons, with the ATG start codon in Exon 1 and the TAG stop codon in exon 9. Exons 2–8 were selected as the conditional knockout (cKO) region and flanked by LoxP sites. Deletion of this region should result in the loss of function of the mouse C11B2 gene. To engineer the targeting vector, homologous arms and the cKO region were generated by PCR using BAC clone RP23-357O1 as a template. Cas9, gRNA and targeting vector were co-injected into fertilized eggs for cKO mouse production. The pups were genotyped by PCR followed by sequencing analysis.

Multiple crosses were conducted between heterozygous C11B2 floxed mice and Pax8-rtTA-LC-1(CRE) mice to produce mice homozygous for the C11B2 floxed allele (C11B2^f/f^) and heterozygous for the Pax8-rtTA/LC-1 transgenes. Unless otherwise specified, all of the mice were bred on a C57BL/6j background. When the mice were 3–4 months of age, renal tubule-specific gene inactivation was induced by administering 2 mg/ml of doxycycline in 5% sucrose drinking water for 14 days. PCR genotyping was used to detect the Pax8-rtTA/LC-1 transgenes and the C11B2 floxed allele. The sequences of primers were as follows: Pax8-rtTA-F, 5’-CCATGTCTAGACTGGACAAGA-3’ and Pax8-rtTA-R, 5’-ATCAATGTATCTTATCATGTCTGG-3’, with a predicted 600-bp product; LC-1-F, 5’-TCGCTGCATTACCGGTCGATGC-3’ and LC-1-R, 5’-CCATGAGTGAACGAACCTGGTCG3’, with a predicted 480-bp product was amplified. The genotyping primers for detecting the C11B2 floxed allele were as follows: C11B2-F1: 5’-ATGCTGATGAGCAGTATTTGGAAC-3’, and C11B2-R1: 5’-GTGCAGTCACCATGCTGAAAATAG-3’, with a predicted 200-bp product from the recombined allele and a 133-bp product from the wild-type (WT) allele. The successful deletion of RT C11B2 was verified by a 437-bp PCR product amplified from renal tissue with the following primer set: C11B2 cKO F: 5’-AACCACTAGCAGGCTGTGCAGG-3’ and C11B2R: 5’-GTGCAGTCACCATGCTGAAAATAG-3’. Tissues without LoxP recombination produced a 630-bp PCR product using the following primers: C11B2 cKO F: 5’-aaccactagcaggctgtgcagg-3’ and C11B2R2: 5’-AAGTGGAATGCCCTCTACCTGAG-3’. To examine the influence of renal knockout of C11B2 gene on C11B2 mRNA expression, we performed RT-PCR with the following primers: C11B2-F, 5’-TGCACCTGGAGCCCTGGGTGGC -3’ and C11B2-R, 5’-GTCATTACCAAGGGGATTGCTG -3’. Both primers are not supposed to bind to CYP11B1 cDNA and yield a 646 bp PCR product amplifying the C11B2 transcript from exons 3–6. As usual, we also used GAPDH to serve as an endogenous control. The primer sequences for GAPDH were as follows: GAPDH-F, 5’-AACTTTGGCATTGTGGAAGG-3’ and GAPDH-R, 5’-GGATGCAGGGATGATGTTCT -3’, which generated a 132-bp PCR product.

### Surgical Procedure

A modified 2K1C murine model of renovascular hypertension was established by placing a polyurethane cuff on the left renal artery, according to a method previously reported with some modifications.^12,13^ We used this method because, according to Lorenz et al.^14^, using conventional U-designed silver clips induces a low success rate of hypertension (40-60%) because the clip laterally presses on the artery, resulting in a constrictions, and the placement of a plastic cuff would result in a constriction in two dimensions (constriction) rather than one (flattening), as with a U-designed silver clip. Due to variability in the levels of hypertension obtained with the conventional U-design silver clip, Lorenz et al.^14^ used rounded polyurethane tubing to initiate renal artery stenosis in mice to avoid these disadvantages. Therefore, we used a small segment of polyurethane tubing (internal diameter: 0.30 mm; outside diameter: 0.63 mm; wall thickness: 0.16 mm; MRE: 025, Braintree Scientific), sliced lengthwise, as a cuff to produce a constriction around the left renal artery.^12^ The tube was cut open lengthwise and placed around the left main renal artery, approximately equidistant between the aorta and renal bifurcation. The cuff was closed and held in place with two sutures. In the 2K1C group, mice that didn’t develop hypertension or the clipped kidney became atrophic were excluded. Sham mice underwent the same procedure, but the cuff was not placed. In order to minimize discomfort, distress, pain, and injury, all animal surgeries were performed during general anesthesia using 2% isoflurane. After the surgical procedure, the supplemental heat was provided, and animals were monitored for respiratory rate and physical activity every 20 minutes for up 3 hours. Buprenorphine-SR was given to control pain (0.01 mg/kg, s.c.) after the surgery.^3^

### Mouse Experiments

Male 12-16-week-old RT C11B2 KO mice, and their respective littermate floxed controls (body weight: roughly 25g) were studied. One week prior to the 2K1C/sham surgery, under anesthesia by 2% isoflurane, the radiotelemetric device was implanted via catheterization of the carotid artery. After collection of baseline blood pressure (BP), all mice were placed in metabolic cage for 24-h basal urine collection. They were then randomly divided to undergo 2K1C operation. Mice were not disturbed during the BP recording period. BP was recorded for 4 hours from 5:00 PM to 9:00 PM every 3 days for 24 days. Following the BP recording, mice were placed in a metabolic cage (MMC100, Hatteras Instruments, Cary, North Carolina) for 24-h urine collection. At the end of the experiment, blood was withdrawn under general anesthesia by puncturing the vena cava. The blood was then centrifuged at 4,000 rpm for 10 minutes to collect the plasma. The animals’ kidneys were harvested and cut into the cortex and the medulla (inner and outer medulla) and snap-frozen^15^.

### Immunoblotting

Renal tissues including the cortex and the inner medulla were lysed and subsequently sonicated in PBS that contained 1% Triton x-100, 250 μM phenylmethanesulfonyl fluoride (PMSF), 2 mM EDTA, and 5 mM dithiothrietol (DTT) (pH 7.5).^16^ Protein concentrations were determined using Coomassie reagent. Forty micrograms of protein for each sample was denatured in boiling water for 10 minutes, separated via SDS-PAGE, and transferred onto nitrocellulose membranes. The blots were blocked for 1 hour with 5% nonfat dry milk in Tris-buffered saline (TBS) and then incubated overnight with primary antibody. After being washed with TBS, the blots were incubated with horseradish peroxidase (HRP)-conjugated secondary antibody and visualized using Enhanced Chemiluminescence (ECL). The blots were quantitated using Image-Pro Plus. The primary antibodies were as follows: rabbit anti-fibronectin antibody (cat no. ab23750; Abcam), rabbit anti-α-smooth muscle actin (α-SMA) antibody (cat no. ab5694; Abcam), rabbit anti-α-ENaC antibody (cat no. SPC-403D; Stressmarq Biosciences), and rabbit anti-GAPDH antibody (cat no. 5174S; Cell Signaling Technology).^17 18^

### qRT-PCR

Total RNA isolation and reverse transcription were performed as previously described.^18,19^ Oligonucleotides were designed using Primer3 software (available at http://frodo.wi.mit.edu/cgi-bin/primer3/primer3_www.cgi). Primers were as follows: for α-ENaC: 5’-gcgacaacaatccccaag-3’ (sense) and 5’-tgaagcgacaggtgaagatg-3’ (antisense); for β-ENaC: 5’-aagcacctgtaatgcccaag-3’ (sense) and 5’-atagcccatccccaccag-3’ (antisense); for γ-ENaC: 5’-cgaagaaactggtgggattt-3’ (sense) and 5’-gatggtggaaaagcgtgaag-3’ (antisense); for IL-1β: 5’-TGCCACCTTTTGACAGTGATG -3’ (sense) and 5’-TGATGTGCTGCTGCGAGATT -3’ (antisense); for TNF-α: 5’-CCCTCACACTCAGATCATCTTCT -3’ (sense) and 5’-GCTACGACGTGGGCTACAG -3’ (antisense); for MCP-1: 5’-TTAAAAACCTGGATCGGAACCAA -3’ (sense) and 5’-GCATTAGCTTCAGATTTACGGGT -3’ (antisense); for TGF-β1: 5’-GGAGAGCCCTGGATACCAAC -3’ (sense) and 5’-CAACCCAGGTCCTTCCTAAA -3’ (antisense); for Fibronectin: 5’-cgaggtgacagagaccacaa-3’ (sense) and 5’-ctggagtcaagccagacaca-3’ (antisense); for alpha-smooth muscle actin (α-SMA): 5’-gaggcaccactgaaccctaa-3’ (sense) and 5’-catctccagagtccagcaca-3’ (antisense); C11B2: 5’-ACCTGCCCTTGCTGAGGGCTGC-3’ (sense), 5’-GCACCCCAAAGCCAAAGGCTAG-3’ (antisense) or 5’-TGCACCTGGAGCCCTG GGTGGC-3’ (sense), 5’-CAAGAAGTCCCTTGCTACCATG-3’ (antisense); and for GAPDH: 5’-gtcttcactaccatggagaagg-3’ (sense) and 5’-tcatggatgaccttggccag-3’ (antisense).^16,20,21^

### Enzyme Immunoassay

Biologic fluids were subjected to ELISA determination of Aldo (catalog no. 501090; Cayman Chemical, Michigan, USA) according to the manufacturers’ instructions. Urine albumin was determined using the DAC Vantage Analyzer (SIEMENS).^15^

### Histopathological Analysis

The kidneys were immersion fixed in 10% buffered formalin, embedded in paraffin, and sectioned into 4-µm-thick slices. Masson’s trichrome staining was used to examine kidney histology and fibrosis. Renal tubulointerstitial indices were assessed using a semiquantitative scoring system, as previously described.^22,23^ In brief, renal fibrosis was determined as blue staining under light microscopy. The area of renal fibrosis was calculated by 10 randomly selected non-overlapping fields at 200×magnification using NIH Image J software (National Institutes of Health, Bethesda, MD). Images were captured by a Leica fluorescence microscope.^15^

### Sodium Nitroprusside (SNP) Experiments

For the sodium nitroprusside (SNP) experiments, male C57BL/6j WT mice aged 7–8 weeks, weighing approximately 18–22 g, were obtained from Jackson Laboratory. The mice were randomly allocated into two groups: 2K1C and 2K1C/SNP. Five days prior to the 2K1C surgery, a radiotelemetry device was implanted under anesthesia via catheterization of the carotid artery. After we recorded baseline BP, all mice underwent the 2K1C surgery. Two weeks after the 2K1C surgery, once the animals’ BP had gradually increased, the mice were randomly and blindly infused with either the vehicle or SNP (1 mg/kg/day, protected from light)^24,25^ via an osmotic mini-pump (Alzet model 1002, Alza) for 28 days. To ensure the efficacy of SNP, we replaced the pump with a new one containing fresh vehicle or SNP on day 14. During the BP recording period, the mice were left undisturbed, and BP was recorded for 4 hours per day from 5:00 PM until 9:00 PM. On day 24 after the vehicle or SNP treatment, the mice were placed in metabolic cages for 24 hours of urine collection for 3 days. At the conclusion of the experiment, blood was collected via puncture of the vena cava under general anesthesia. The animals’ kidneys were harvested and dissected into the cortex and medulla and then snap-frozen for further analysis.

### Statistics

We used GraphPad Prism software (version 8.4) for the data analysis. The data were summarized as means ± SEM. All data points represent animals that were included in the statistical analyses. Sample sizes were determined on the basis of similar previous studies or pilot experiments. For the BP experiment, statistical significance was determined using two-way ANOVA with repeated measurements. The other animal experiments were performed using regular two-way ANOVA with the Bonferroni test for multiple comparisons or the unpaired two-tailed Student’s t test for two comparisons. A *P* value less than 0.05 was considered to indicate statistical significance. ^17,26^

## Results

### Generation of RT C11B2 KO Mice

We used CRISPR/Cas9 to insert two LoxP sites into introns 1 and 8 of the C11B2 gene (Fig. 1A). As shown in Fig. 1B, the genotype of C11B2 floxed allele was analyzed by PCR. C11B2^f/f^ mice developed normally and were fertile. Multiple crosses were conducted between heterozygous C11B2 floxed mice and Pax8-rtTA-LC-1(Cre) mice to produce mice homozygous for C11B2^f/f^ and heterozygous for the Pax8-rtTA/LC-1 transgenes. When the mice were 3–4 months of age, renal tubule-specific gene inactivation was induced by administering 2 mg/ml of doxycycline in 5% sucrose drinking water for 14 days; the resulting animals were termed RT C11B2 KO mice. As shown in Fig. 1C, DNA recombination in RT C11B2 KO mice was detected only in the kidneys but not in other tissues. In agreement with this finding, C11B2 mRNA, as assessed by RT-PCR, was detected in the renal cortex and medulla of floxed mice but was absent in the RT C11B2 KO mice (Fig. 1D). On the other hand, adrenal C11B2 mRNA did not differ between the two genotypes (Fig. 1D). RT C11B2 KO mice excreted 58% less urinary compared with the C11B2 floxed mice under basal conditions (Fig. 1E). More importantly, clipping-induced urinary Aldo was reduced by 78% (Fig. 1E), and plasma Aldo remained unchanged (Fig. 1F). RT C11B2 KO mice exhibited the same normal phenotype under basal condition as C11B2^f/f^ mice. We accordingly established a novel mouse line of renal tubule-specific deletion of C11B2 that offers an unique opportunity to investigate the function of intrarenal generation of Aldo.

**Fig. 1.**
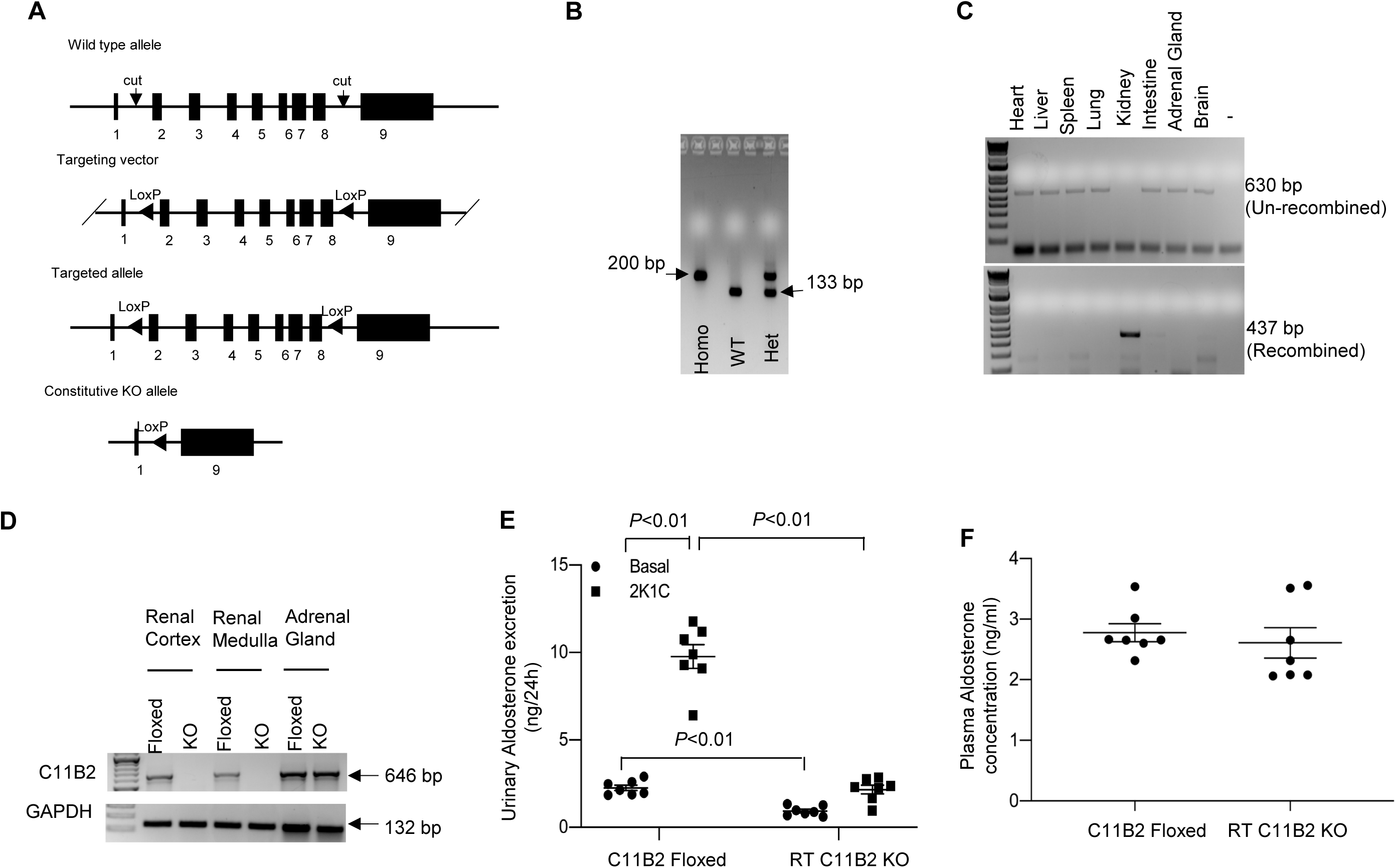
Generation and characterization of renal tubule-specific C11B2 KO mice (RT C11B2 KO). (A) Schematic illustration of the strategy for generating the C11B2 floxed allele using the CRISP/Cas9 method. Briefly, exons 2–8 were selected as a cKO region and flanked by LoxP sites. To engineer the targeting vector, homologous arms and cKO region were generated by PCR using BAC clone RP23-357O1 as a template. Cas9, gRNA, and the targeting vector were co-injected into fertilized eggs for cKO mouse production. (B) C11B2 floxed mice were genotyped by PCR showing the detection of PCR products from the WT or floxed allele. (C) DNA recombination in male 3–4-month-old RT C11B2 KO mice was examined by PCR using PCR primers flanking the two LoxP sites. The 630- and 437-bp products represent the un-recombined and recombined alleles, respectively. DNA recombination was specifically detected in the kidneys but not other tissues. (D) RT-PCR analysis of C11B2 mRNA expression in the renal cortex and medulla versus adrenal glands of male 3–4-month-old C11B2^f/f^ and RT C11B2 KO mice. Agarose gel electrophoresis showed that the 646-bp PCR products were detected in the renal cortex and medulla of floxed but not RT C11B2 KO mice. On the other hand, adrenal CY11P2 mRNA expression did not differ between the genotypes. (E) Urinary excretion of Aldo in male 3–4-month-old C11B2 floxed and RT CYP11B2 KO mice under basal conditions and 4 weeks after the 2K1C procedure. (F) at the end of the experiment, blood was withdrawn from vena cava under anesthesia. Aldo in the urine and plasma was measured using ELISA. Data are means ± SE. n = 7 per group.

### Ablation of C11B2 in RT Attenuates 2K1C-induced Renovascular Hypertension

We examined the BP phenotype of RT C11B2 KO mice during 2K1C-induced renovascular hypertension. Radiotelemetry was used to monitor changes in mean arterial pressure (MAP), systolic blood pressure (SBP), diastolic blood pressure (DBP), and heart rate (HR) in RT C11B2 KO mice and C11B2^f/f^ mice following the 2K1C procedure. The 2K1C procedure gradually elevated MAP, SBP, and DBP; but decreased HR. These effects were observed in both genotypes, but the hypertensive and bradycardia responses to the 2K1C procedure were less pronounced in the RT C11B2 KO mice compared with the C11B2^f/f^ controls (Fig. 2). These results suggest that activation of RT C11B2 partially contributes to the hypertensive response to the 2K1C procedure.

**Fig. 2.**
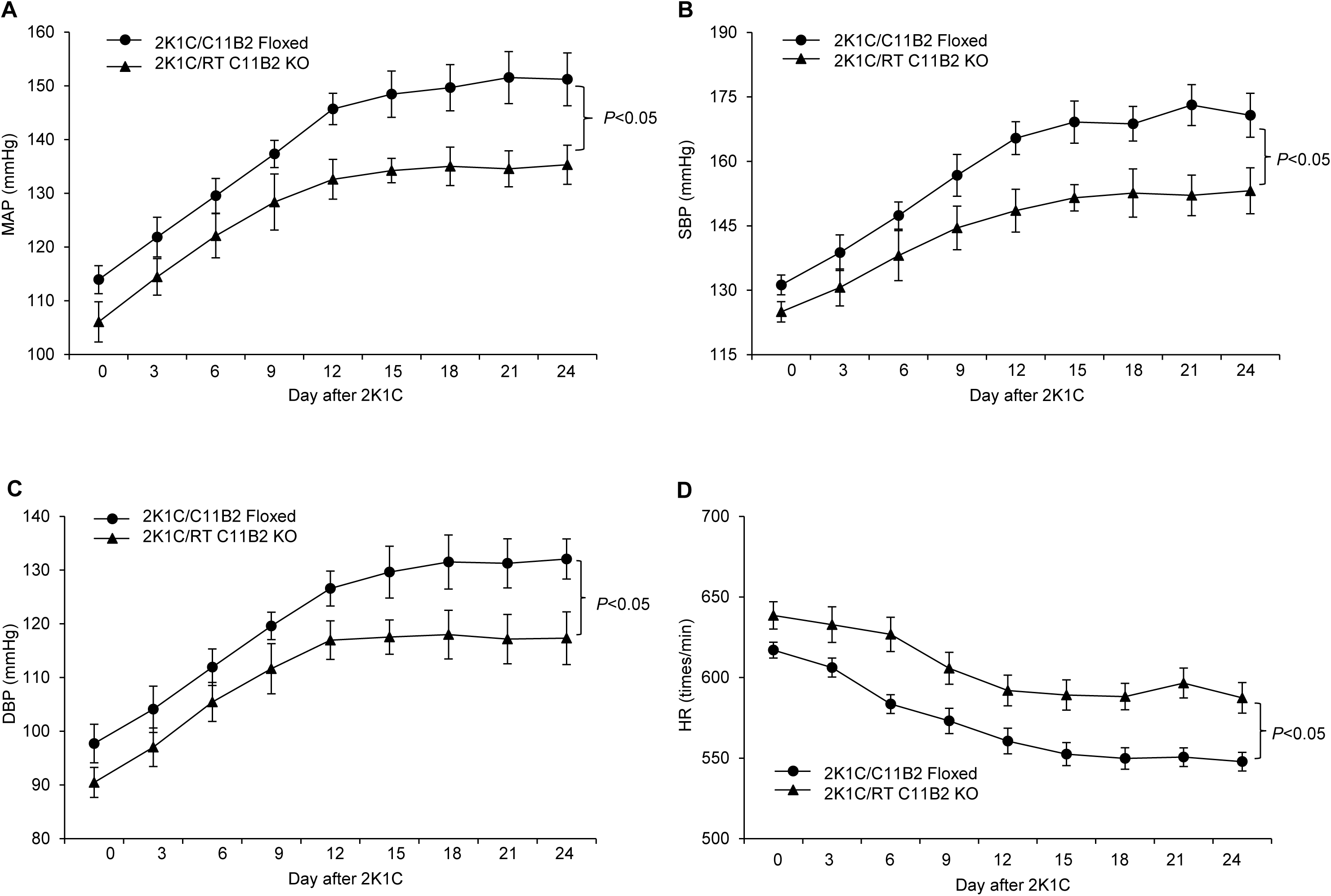
The role of RT C11B2 in 2K1C-induced renovascular hypertension. Following the 2K1C procedure, BP was monitored over the course of four hours from 5:00 PM to 9:00 PM using radiotelemetry. (A) Mean arterial pressure (MAP). (B) Systolic blood pressure (SBP). (C) Diastolic blood pressure (DBP). (D) Heart rate (HR). Statistical significance was determined by using two-way ANOVA with repeated measurements, and *P* values are indicated in the figure. C11B2 floxed/2K1C: n = 7; RT C11B2 KO/2K1C: n = 7. Data are means ± SE.

### Assessment of Renal Injury

The 2K1C model is well known to feature renal pathologies, including inflammation, fibrosis, and hypertension. ^4,27^ Therefore, we examined markers of renal fibrosis such as the expression of fibronectin and α-SMA and urinary excretion of albumin, KIM-1, and NGAL 4 weeks after the 2K1C procedure. RT C11B2 KO mice exhibited a significant attention of clipping-induced renal fibrosis, as reflected by changes in renal protein abundances of fibronectin and α-SMA, compared with the floxed controls (Fig. 3A). Moreover, urinary excretion of albumin, KIM-1, and NGAL were increased in C11B2 floxed mice; excretion was attenuated in RT C11B2 KO mice (Fig. 3B-D). Additionally, we performed Masson Trichrome staining to evaluate renal fibrosis. As demonstrated by the semiquantitative tubulointerstitial fibrosis index of the kidney sections (Fig. 3E), 2K1C-treated C11B2 floxed mice exhibited increased accumulation of the extracellular matrix in the clipped kidney, which was attenuated in RT C11B2 KO mice (Fig. 3F). On the other hand, Masson Trichrome staining revealed no significant tubulointerstitial fibrosis in the contralateral kidneys of 2K1C mice, irrespective of genotype (Fig. 3F). Consistent with this observation, clipping induced parallel increases in renal mRNA expression of fibronectin, α-SMA, IL-1β, TNF-α, MCP-1, and TGF-β1 in C11B2 floxed mice, which were all blunted in RT C11B2 KO mice (Fig. 4). Together, these results suggest a pathogenic role of intrarenal Aldo in clipping-induced renal injury.

**Fig. 3.**
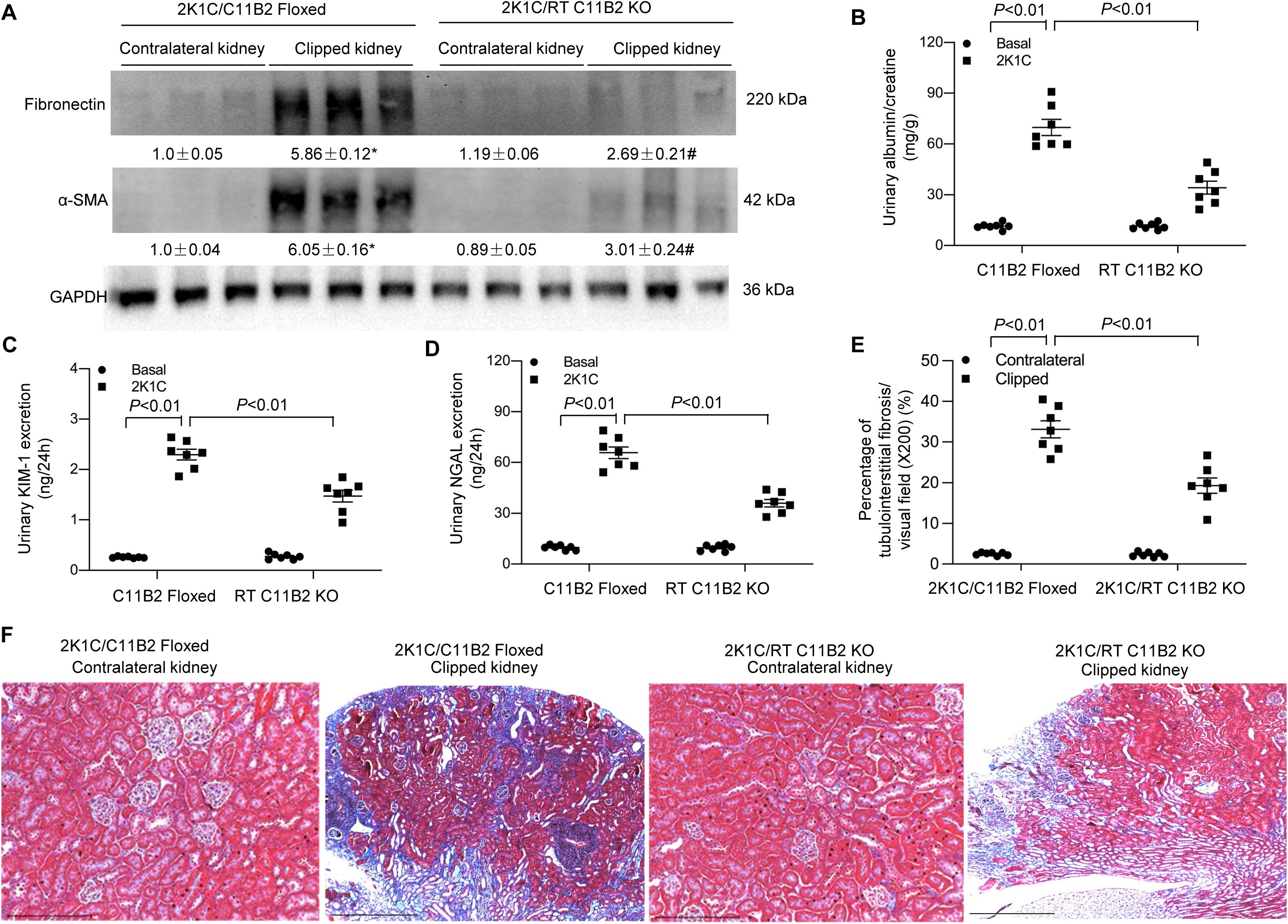
Role of RT C11B2 in 2K1C-induced ischemic nephropathy. (A) The renal cortex of contralateral and clipped kidneys of C11B2^f/f^ and RT C11B2 KO mice was subjected to immunoblotting analysis of protein expression of fibronectin and α-SMA. Protein abundances were analyzed using densitometry and normalized by GAPDH. The values are shown underneath the representative blots. The urinary albumin-to-creatinine ratio (B), urinary excretion of KIM-1 (C) and NGAL (D) were determined under basal conditions and 4 weeks after 2K1C. (E) The percentage of tubulointerstitial fibrosis/visual field (%). (F) Representative micrographs from Masson Trichrome staining. Statistical significance was determined using two-way ANOVA with the Bonferroni test for multiple comparisons. *, *P*<0.01 vs. contralateral kidney/C11B2^f/f^. #, *P*<0.01 vs. clipped kidney/C11B2^f/f^ In panels B-E, *P* values are indicated. C11B2 floxed/2K1C: n = 7; RT C11B2 KO/2K1C: n = 7. Data are means ± SE.

**Fig. 4.**
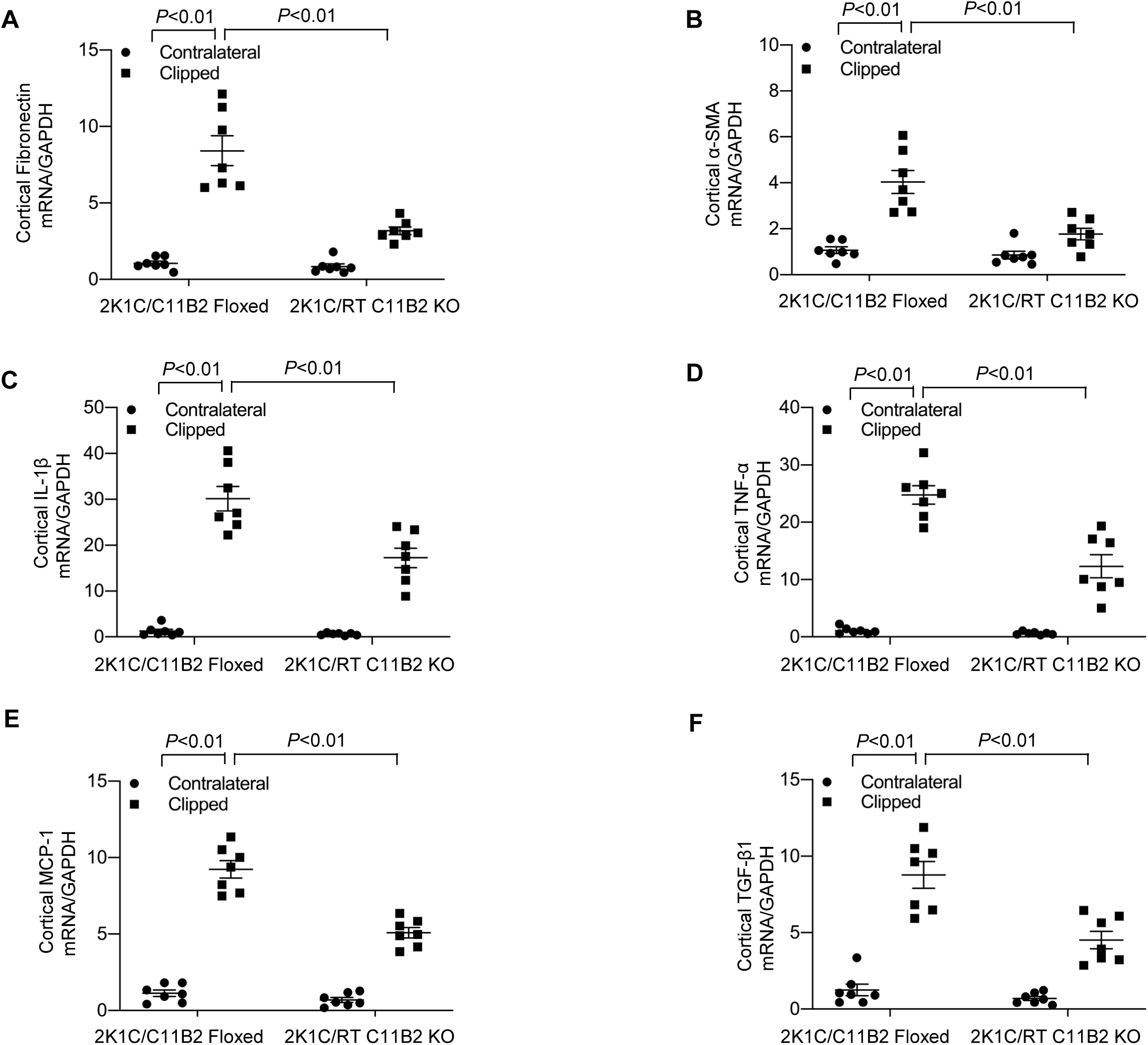
Assessment of renal injury markers. The renal cortex of contralateral and clipped kidneys were subjected to qRT-PCR analysis of mRNA expression of fibronectin (A), α-SMA (B), IL-1β (C), TNF-α (D), MCP-1 (E), and TGF-β1 (F) normalized by GAPDH. Statistical significance was determined using two-way ANOVA with the Bonferroni test for multiple comparisons. *P* values are indicated in the figure. C11B2 floxed/2K1C: n = 7; RT C11B2 KO/2K1C: n = 7. Data are means ± SE.

### Assessment of Renal ENaC

The enhancement of ENaC-mediated Na^+^ reabsorption in the distal nephron plays an important role in the pathogenesis of hypertension. ^28,29^ Of the three subunits of ENaC, the α subunit is the rate-limiting component in ENaC-mediated Na^+^ transport and is sensitive to Aldo. ^30^ Moreover, renal medullary α-ENaC is specifically targeted by sPRR to induce hypertension during overactivation of the RAS. ^31^ We therefore examined mRNA expression of the three ENaC subunits in the renal cortex and medulla of our experimental models using qRT-PCR. Clipping-induced renal medullary α-ENaC mRNA expression was significantly decreased in RT C11B2 KO mice compared with the C11B2 floxed controls (Fig. 5A). On the other hand, renal cortical α-ENaC mRNA expression remained unaffected in RT C11B2 KO mice (Fig. 5B). Moreover, renal mRNA expression of the β-ENaC and γ-ENaC subunits remained constant across various experimental groups, irrespective of the genotype or kidney regions (Fig. 5A-B). These results collectively support renal medullary α-ENaC as a target of intrarenal Aldo during renovascular hypertension.

**Fig. 5.**
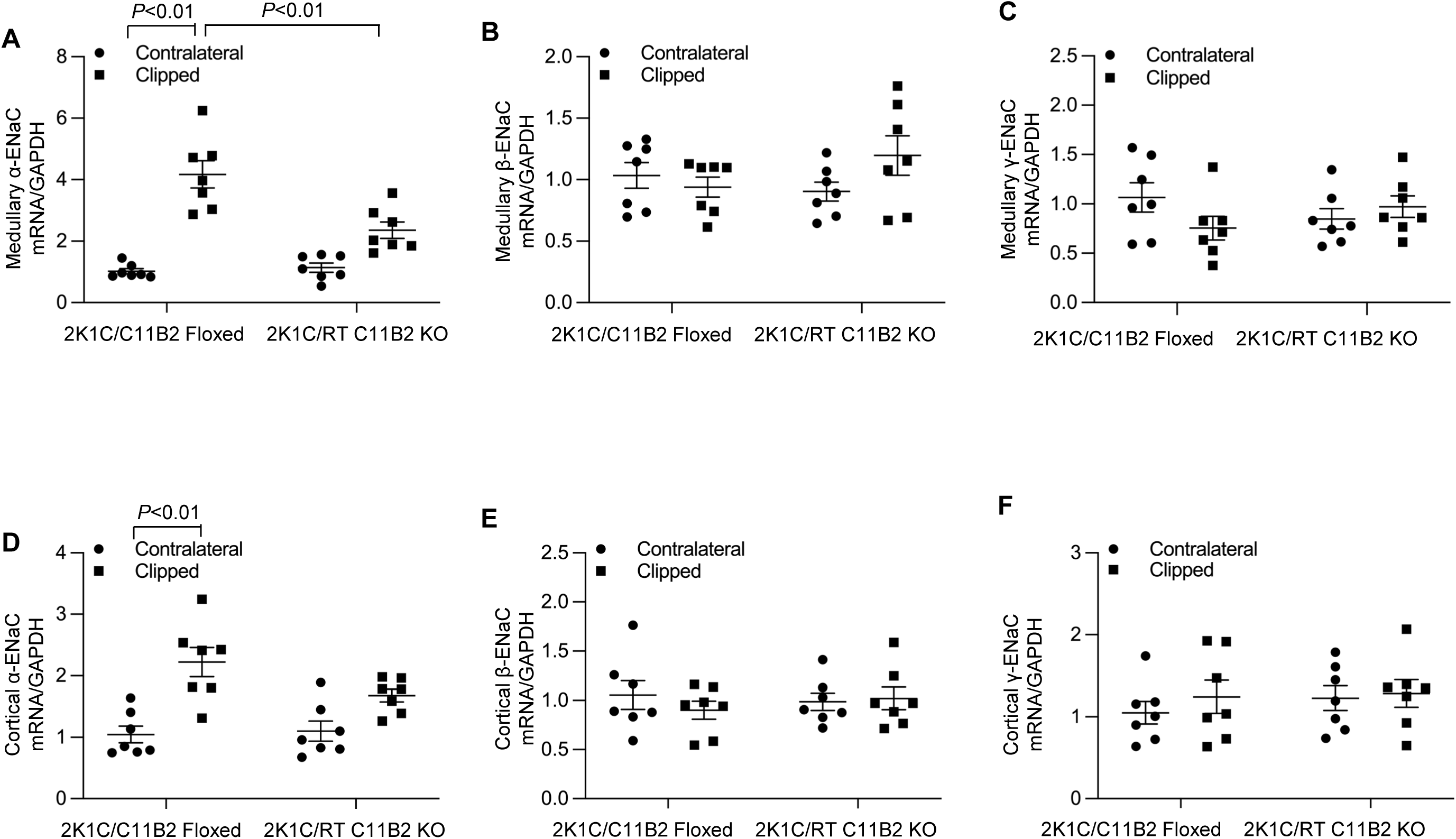
Assessment of renal regional expression of ENaC subunits. The mRNA expression of the three subunits of ENaC (α-ENaC in renal medulla (A-C) and cortex (D-F) of the contralateral and clipped kidneys in C11B2 floxed and RT C11B2 KO mice was determined at 4 weeks after 2K1C by qRT-PCR and normalized by GAPDH. (A, D) α-ENaC. (B, E) β-ENaC. (C-F) γ-ENaC. Statistical significance was determined using two-way ANOVA with the Bonferroni test for multiple comparisons or the unpaired two-tailed Student’s T test for two comparisons. *P* values are indicated in the figure. C11B2^f/f^/2K1C: n = 7; RT C11B2 KO/2K1C: n = 7. Data are means ± SE.

### The Phenotypes of RT C11B2 KO Mice in an Acute 2K1C Model

Although the 2K1C model is widely used as a chronic model of renovascular hypertension and renal fibrosis, we assessed markers of acute kidney injury (AKI) 24 hours after 2K1C in the absence of elevated BP. It typically takes roughly 1 week to observe a noticeable elevation in BP following the 2K1C procedure. ^3^ Twenty-four hours after the 2K1C procedure, urinary excretion of albumin, KIM-1, and NGAL increased in the C11B2 floxed mice, which is indicative of AKI (Fig. 6A-C). Urinary excretion of albumin, KIM-1, and NGAL decreased in RT C11B2 KO mice (Fig. 6A-C). Additionally, urinary Aldo excretion was elevated by 1-day clipping in C11B2 floxed mice; this increase was significantly muted in RT C11B2 KO mice (Fig. 6D). These results support the concept that intrarenal Aldo mediates clipping-induced renal injury, independent of hypertension.

**Fig. 6.**
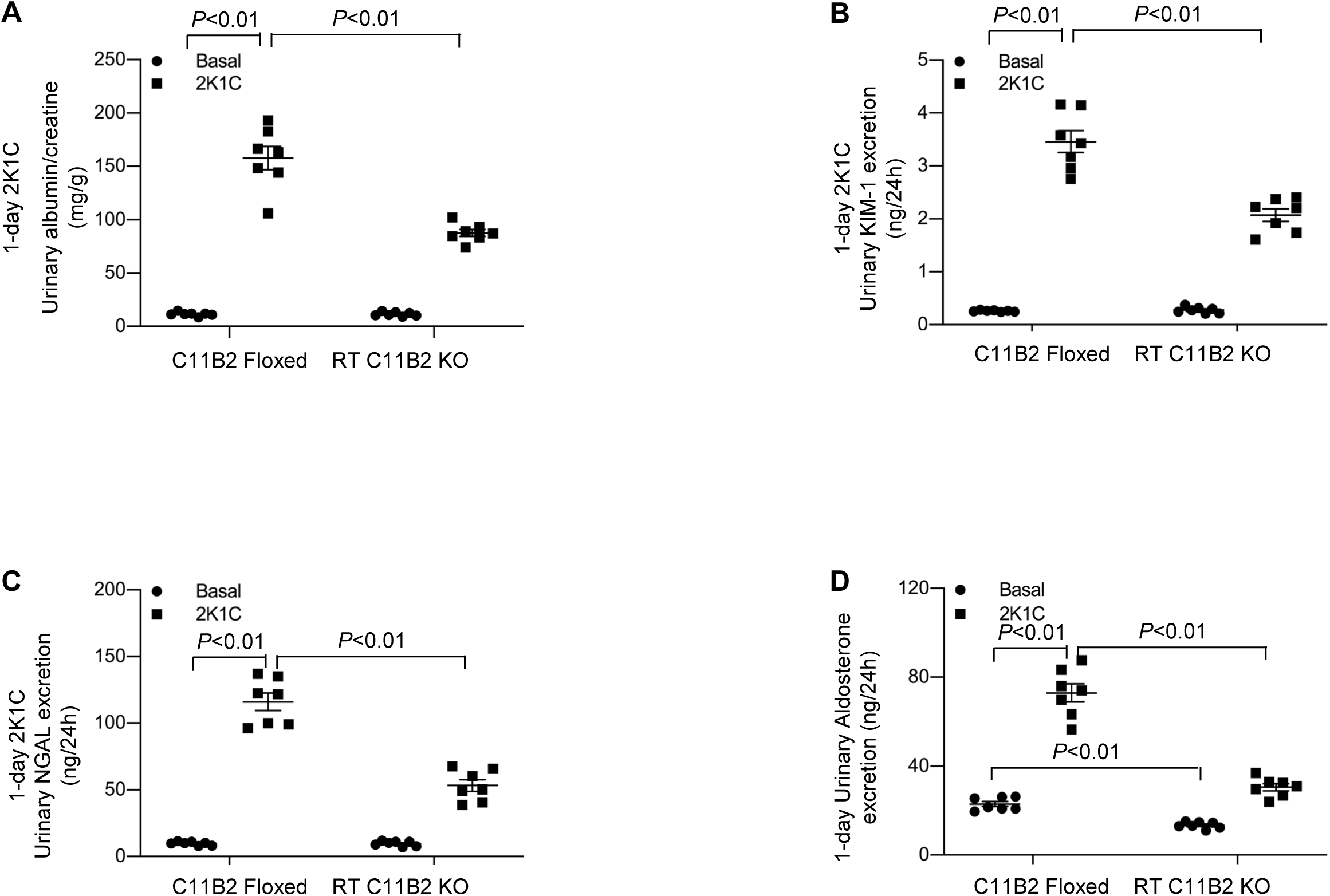
Clipping-induced acute kidney injury in RT C11B2 KO mice. The urinary albumin-to-creatinine ratio (A), KIM-1 (B), NGAL (C), and Aldo excretion (D) were detected under basal condition and 24 hours after 2K1C. Statistical significance was determined using two-way ANOVA with the Bonferroni test for multiple comparisons, and *P* values are indicated in the figure. C11B2 floxed/2K1C: n = 7; RT C11B2 KO/2K1C: n = 7. Data are means ± SE.

### A Lack of Effect of Non-RAAS Antihypertensive Therapy on Renal Injury in a 2K1C Model

To further clarify the relationship between hypertension and renal injury in the 2K1C model, we examined the effect of SNP, a commonly used non-RAAS vasodilator, on renal injury in C57/BL6/j mice subjected to the 2K1C procedure. Interestingly, while SNP significantly and rapidly attenuated the hypertensive response induced by the 2K1C procedure (Fig. 7A-D), it did not affect albuminuria (Fig. 7E). Consistent with this observation, clipping-induced elevations of mRNA levels of fibronectin and α-SMA were also unaffected (Fig. 7F-G). These results provide additional evidence that clipping-induced renal injury may not be a direct consequence of hypertension in the 2K1C model.

**Fig. 7.**
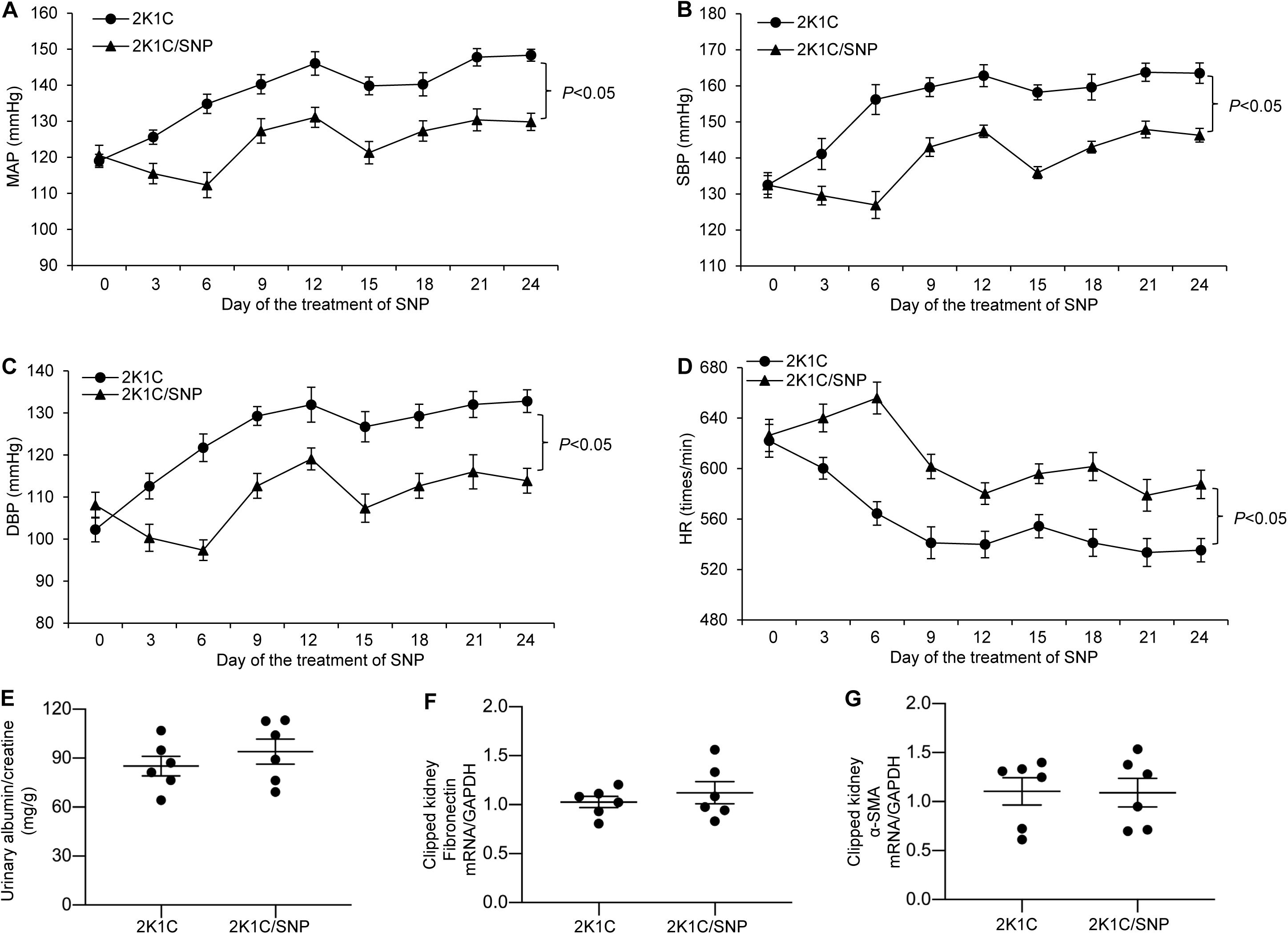
Effects of SNP on renal injury in the 2K1C model. Under anesthesia, male 3–4-month old C57BL/6j mice were instrumented with radiotelemetric devices. After five days of recovery, the mice underwent the 2K1C procedure. They were then randomly divided to receive subcutaneous infusion of vehicle or SNP at 1 mg/kg/day via an osmotic minipump for 4 weeks. Blood pressure was collected at the indicated time points from 5:00 PM to 9:00 PM using radiotelemetry. (A) MAP. (B) SBP. (C) DBP. (D) HR. Statistical significance was determined using two-way ANOVA with repeated measurements, and the *P* values are indicated in the figure. The urinary albumin-to-creatinine ratio (E) was detected 4 weeks post-SNP. The renal cortex of clipped kidneys was subjected to qRT-PCR analysis of mRNA expression of fibronectin (F) and α-SMA (G). mRNA expression was normalized by GAPDH. Statistical significance was determined by using the unpaired two-tailed Student’s T test for two comparisons. 2K1C: n = 6; 2K1C/SNP: n = 6. Data are means ± SE.

### The Regulation of Intrarenal Aldo Synthesis in CD PRR/renin KO Mice Following the 2K1C Procedure

Our research group recently showed that CD-specific deletion of PRR or renin induced a partial attenuation of hypertension and a more significant improvement in ischemic nephropathy induced by the 2K1C procedure^3^. This phenotype is nearly analogous to that of RT C11B2 KO mice. It appears likely that intrarenal Aldo generation may serve as an integrative component of an intact PRR-dependent intrarenal RAAS. To define CD PRR/renin as the upstream regulators of intrarenal Aldo generation, we examined renal C11B2 expression and Aldo production in CD PRR KO and CD renin KO mice subjected to the 2K1C procedure. Following the 2K1C surgery, C11B2 mRNA expression was upregulated in both the renal cortex and medulla of the clipped kidneys in PRR floxed mice. Interestingly, CD PRR KO selectively blunted clipping-induced C11B2 mRNA expression in the renal medulla, but not in the cortex (Fig. 8A-B). Urinary excretion and the plasma concentrations of Aldo were both elevated in PRR floxed mice after the 2K1C procedure; however, CD PRR KO only attenuated the increases in urinary Aldo excretion—plasma levels remained unchanged (Fig. 8C-D). Similarly, 2K1C-induced increases in renal medullary C11B2 expression and urinary Aldo excretion were both blunted in CD renin KO mice. That finding contrasts to the static responses of renal cortical C11B2 expression and plasma Aldo levels (Fig. 8E-H). Together, these results provide a strong link between CD PRR/renin and renal medullary Aldo biosynthesis in the 2K1C model.

**Fig. 8.**
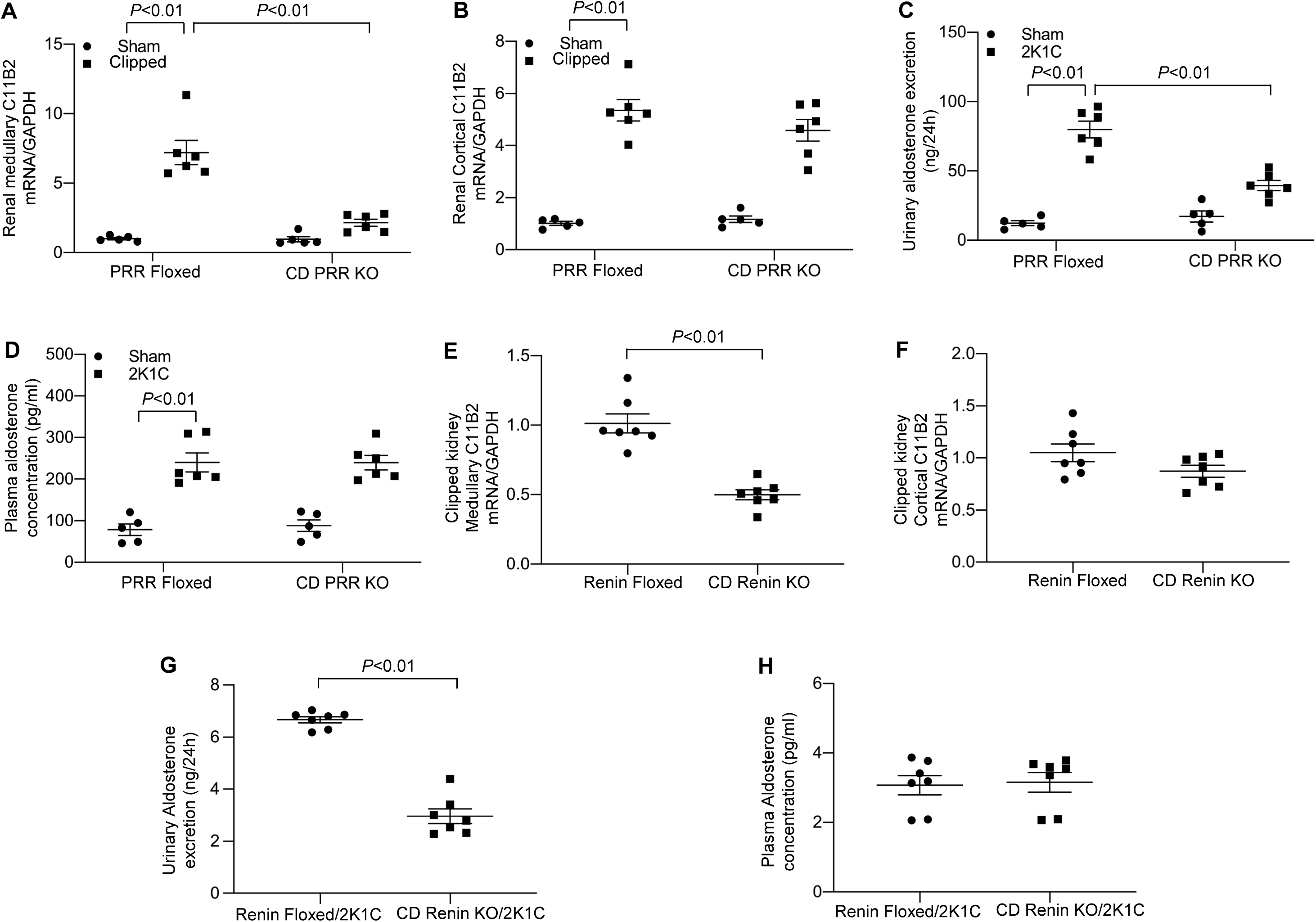
The role of CD PRR/renin in the regulation of intramedullary Aldo biosynthesis during renal artery clipping. Under anesthesia, male 3–4-month-old CD PRR KO and CY11B2^f/f^ mice underwent a sham operation or the 2K1C procedure. Four weeks later, the renal medulla (A) and cortex (B) were subjected to qRT-PCR analysis of C11B2 mRNA expression normalized by GAPDH. Urinary excretion (C) and plasma concentrations (D) of Aldo were determined by ELISA. In a separate experiment, male 3–4-month-old CD renin KO and Renin^f/f^ mice underwent the 2K1C procedure and were analyzed 4 weeks later. C11B2 mRNA expression of the renal medulla (E) and cortex (F) of clipped kidneys in Renin^f/f^ or CD renin KO mice was determined by qRT-PCR. Urinary excretion and plasma concentrations of Aldo were analyzed by ELISA. Renin^f/f^/2K1C: n = 7; CD renin KO/2K1C: n = 7. Statistical significance was determined using two-way ANOVA with the Bonferroni test for multiple comparisons or the unpaired two-tailed Student’s T test for two comparisons. *P* values are indicated in the figure. Data are means ± SE.

## Discussion

This study is the first to provide genetic evidence establishing an essential role of intrarenal Aldo in the 2K1C model. RT C11B2 KO partially attenuated hypertension but more significantly improved renal information and fibrosis. During the 24 hours after the 2K1C procedure, albuminuria was attenuated in KO mice in the absence of hypertension. Additionally, decreasing BP with a NO donor SNP did not affect clipping-induced renal injury in WT mice. Likely, intrarenal Aldo plays a primary role in ischemic nephropathy but a secondary role in hypertension development in the 2K1C model.

The successful generation of a RT C11B2 KO model in the present study is evidenced by PCR-based confirmation of renal-specific DNA recombination of the C11B2 gene, the disappearance of renal C11B2 mRNA, and the significant reduction in urinary—but not plasma—Aldo. Consistent with these findings, adrenal C11B2 expression was unaffected. It seems reasonable to speculate that RT C11B2 KO attenuates 2K1C-induced renovascular hypertension and ischemic nephropathy by decreasing the generation of intrarenal Aldo. Intrarenal Aldo elicits a hypertensive response via activation of the renal medullary α-ENaC; its renal pathogenic role likely depends on the well-described pro-inflammatory and pro-fibrotic action of Aldo. These results strongly support the concept that intrarenal generation of Aldo derives from CD PRR/renin-dependent renal medullary C11B2 expression, which represents a key contributor to hypertension and renal injury in the 2K1C model. Moreover, decreasing BP with a NO donor did not affect renal injury in 2K1C C57/BL6 mice. Intrarenal RAAS seems to play a primary role in ischemic nephropathy but a secondary role in hypertension development in the 2K1C model.

The direct role of intrarenal RAAS in ischemic nephropathy was further tested during 24 hours of the 2K1C procedure. In a chronic 2K1C model, BP typically starts to increase after several days but not within the first 24 hours; it reaches a plateau within 1–2 weeks. Therefore, the acute 2K1C model can be designed to be free of the confounding influence of hypertension. Using this protocol, we were able to demonstrate that clipping-induced acute ischemic nephropathy was significantly attenuated in RT C11B2 KO mice, an effect likely independent of hypertension. Similarly, contralateral kidneys in the chronic 2K1C models showed no obvious tubulointerstitial fibrosis. Moreover, administration of a NO donor SNP effectively decreased 2K1C-induced hypertension. On the other hand, this antihypertensive effect was not associated with improved indices of renal injury. Together, these results support the idea that ischemia-induced renal injury in the 2K1C model is likely the cause, rather than the consequence, of hypertension. These results may challenge the view of nephropathy as the consequence of hypertension in the 2K1C model. Indeed, although hypertension is widely considered to be one of the leading causes of CKD after diabetes, the causality of hypertension and CKD has not been firmly established. No prospective controlled clinical trials have been conducted in primary hypertension patients with renal events as primary endpoints to date.^32^ Moreover, the Systolic Blood Pressure Intervention Trial (SPRINT) demonstrated that decreasing blood pressure below current targets provides additional cardiovascular and mortality benefits but induces an acute decrease in eGFR, which is associated with an increased risk of CKD.^33^ Our study calls for further basic and clinical investigation of the precise role of hypertension in the pathogenesis of kidney disease, particularly during renal artery stenosis.

Our group has consistently demonstrated that pharmacological antagonism of renal medullary PRR,^34^ CD-specific deletion of PRR^3,35^ and mutagenesis of the cleavage site of PRR^15,36^ selectively blunted renal medullary expression of α-ENaC but not the β- or γ-subunits. Indeed, among the three subunits, the synthesis of α-ENaC is a limiting factor in the assembly of the ENaC complex.^37^ In agreement with these findings, the present study showed that both renal cortical and renal medullary expression of α-ENaC were all elevated in the clipped kidney of C11B2 floxed mice, but only the renal medullary expression of α-ENaC was blunted in the RT C11B2 KO mice. Furthermore, a selective increase in α-ENaC abundance is usually an indicator of elevated Aldo,^30,38^ Our findings suggest that the selective increase in medullary α-ENaC abundance is a result of heightened intrarenal Aldo levels. It is reasonable to speculate that CD PRR/renin-dependent activation of intrarenal Aldo contributes to the upregulation of renal medullary α-ENaC, which promotes 2K1C-induced hypertension via its sodium-retaining action in the distal nephron.

We would like to acknowledge several limitations to this study. For example, due to the lack of a specific anti-C11B2 antibody, we were unable to investigate the intrarenal distribution of C11B2. A homology of over 90% of cDNA sequences between C11B2 and C11B1 also hinders the development of a C11B2-specific RNA probe for determining the intrarenal distribution of C11B2 mRNA. For these reasons, the deletion of C11B2 in the present study was conducted in the entire renal tubule rather than a specific nephron segment. Although the renal medulla appears to be a major site of local Aldo synthesis under the control of CD PRR/renin, the specific renal medullary cell type capable of synthesizing Aldo remains unknown. Additionally, a human study was not conducted to investigate the clinical relevance of intrarenal Aldo in patients with TRH, renovascular hypertension, or ischemic nephropathy. When a C11B2-specfic antibody becomes available in the future, human kidneys can be analyzed for C11B2 protein expression.

In summary, this work defined the function and regulation of intrarenal Aldo biosynthesis in the 2K1C model. Renal tubule-specific deletion of C11B2 remarkably improves renal fibrosis and inflammation, and partially attenuated hypertension in this model. We have provided further evidence in support of CD PRR as a regulator of renal medullary C11B2. Overall, these findings highlight the crucial role of intrarenal Aldo biosynthesis as a key component of the intrarenal RAAS implicated in the pathogenesis of ischemic nephropathy and renovascular hypertension.

## Data Sharing Statement

The data about detailed protocols, methods, and other useful materials and resources supporting the findings of this study are openly available in repository [figshare] at https://doi.org/10.6084/m9.figshare.24239452.v1. In addition to primary research information, we also shared the critical raw data related to this study.

## Acknowledgments

This work was supported by Veterans Affairs (VA) Merit Review I01BX004871 and Senior Research Career Scientist award IK6BX005223 from the Department of VA, and National Institutes of Health GrantsDK104072 and HL160020.

## Novelty and Significance

### What Is Known?

- Aldo synthase inhibitors have entered a late-stage of clinical development and hold great promise to manage TRH and kidney disease.
- The adrenal glands are believed to be the predominant source of Aldo but the importance of extra-adrenal production of this hormone has largely been ignored.
- The concept and importance of intrarenal RAS are well recognized but local generation of Aldo is often omitted from this local system.
- Despite the well-recognized benefits, anti-RAAS therapies are limited by class toxicities, e.g. hyperkalemia and AKI. Adrenal-derived Aldo plays a key role in the regulation of K^+^ homeostasis.
- Increasing evidence supports a pivotal role of intrarenal RAS in kidney disease and hypertension. However, the function of intrarenal Aldo largely remains elusive.

### What New Information Does This Article Contribute?

- We generated a novel mouse model of renal-specific deletion of C11B2, which offers a powerful tool to dissect the function of intrarenal Aldo without the confounding influence of adrenal-derived Aldo.
- We employed this model to establish an essential role of intrarenal Aldo in ischemic nephropathy and a relatively modest role of this pathway in hypertension development during renal artery constriction.
- Ischemic nephropathy is not merely a consequence of hypertension at least in the 2K1C model.
- Renal medullary Aldo biosynthesis is under the control of the CD PRR/renin axis.
- Intrarenal Aldo promotes pro-inflammatory and pro-fibrotic responses to induce ischemic nephropathy and targets renal medullary α-ENaC to increase Na+ reabsorption and thus hypertension during renal artery constriction.
- These results support the idea that a site-specific inhibition of Aldo synthesis in the renal medulla may represent a more effective and safer Aldo-targeted therapy for management of TRH and kidney disease, particularly those caused by renal artery stenosis.

